# GIN-TONIC: Non-hierarchical full-text indexing for graph-genomes

**DOI:** 10.1101/2023.11.01.565214

**Authors:** Ünsal Öztürk, Marco Mattavelli, Paolo Ribeca

## Abstract

This paper presents a new data structure, GIN-TONIC, designed to index arbitrary string-labelled directed graphs representing, for instance, pangenomes or transcriptomes. GIN-TONIC provides several capabilities not offered by other graph-indexing methods based on the FM-index. It is non-hierarchical, handling a graph as a single monolithic object; it indexes at nucleotide resolution all possible walks in the graph without the need to explicitly store them; it supports exact substring queries in polynomial time and space for all possible walk roots in the graph, even if there are exponentially many walks corresponding to such roots. Specific ad-hoc optimisations, such as a precomputed cache, allow GIN-TONIC to achieve excellent performance for input graphs of various topologies and sizes. Robust scalability capabilities and a querying performance close to that of a linear FM-Index are demonstrated for two real-world applications, a human pangenome and transcriptome. Source code and associated benchmarks are available on GitHub.

**Availability and implementation:** GIN-TONIC and all related programs are available at https://github.com/uensalo/gin.

## 1. Introduction

The importance of being able to represent DNA sequences as complex graphs rather than linear text is well acknowledged in the literature, with increasing attention being given to pangenome-based applications (1; 2; 3; 4). Another relevant use case is transcriptomics, where splicing induces graphs with millions of edges (5).

A data structure commonly used in bioinformatics is the FM-Index (6), which is especially efficient at matching queries against large reference sequences (7; 8; 9). Although implementations of the FM-Index abound, each one striking a different balance between compression and speed (10; 11), a common use for them is to compute the locations of the exact matches of shorter k-mers from a query sequence. This process, which is called seeding, is a common procedure in alignment software (12; 13). However, naïve FM-Index approaches are unable to handle graph genomes.

Aligning reads to graph genomes and pangenomes offers a more comprehensive view of genomic diversity. Algorithmically speaking, however, the central challenge here lies in efficiently indexing these graph structures — unlike linear sequences, graphs can contain complex looping structures and multiple alternative walks, making indexing and alignment more difficult (1; 14) due to the exponential nature of the number of graph walks to be explored (15; 16). An ideal, efficient graph data structure should keep the same good qualities of the FM-Index — notably, the ability to return all the exact matches in polynomial time — while also being able to transparently explore all matches along all possible walks in the graph.

Recent work in graph indexing based on the Burrows-Wheeler transform and the FM-Index includes the vg toolkit (17) and its extensions, which employ the BWT (18), and other extensions of the PBWT (19) for pangenomic-level haplotype indexing (GBWT, gPBWT) (20; 21). Further developments add synthetic haplotypes or greedy graph covers to allow nucleotide-domain indexing (22). HISAT 2 and 3 (23; 24) use a different, hierarchical approach consisting of a global FM-Index for the whole genome and local FM-Indices constructed over smaller genomic regions. In the domain of nucleotide-level indexing, however, neither of these proposals offers guarantees against very complex graphs, meaning that the worst-case scenario might generate very large indices and/or incur very long querying times. Theoretical approaches to fully generalize the combinatorial properties of the Burrows-Wheeler transform to special subclasses of character-labelled graphs have also been proposed. The concept of Wheeler graphs (25) extends the main properties of LF-mapping traversals, establishing properties analogous to those of the Burrows-Wheeler transform, such as the query time being proportional to query length, under the assumption of path coherence and monotonicity. Strategies such as r-indexing (26), tunneling techniques (27), and prefix-free parsing (28), extend the notion of Wheeler graphs to improve performance. Further generalizations are made in (29), in which the authors describe a family of transformations that support matching in FM-Index-like time. A different approach is followed in (30; 31; 32) by considering a relaxation of Wheeler orders to co-lex orders or arbitrary relations, which allows pattern matching on the graph in polynomial time (33). However, scaling to billions of character-labelled nodes remains an algorithmic challenge, rendering algorithmic frameworks based on Wheeler graphs impractical for human-scale pangenomics (15; 16).

Here we introduce a novel data structure called GIN-TONIC (**G**raph **IN**dexing **T**hrough **O**ptimal **N**ear **I**nterval **C**ompaction). It is designed to handle string-labelled directed graphs of arbitrary topology by indexing all possible string walks without explicitly storing them. Crucially, it allows for efficient exact lookups of substring queries of unrestricted length in polynomial time and space (excluding the size of the output, which may be exponential); it does not require the construction of multiple indices or explicit enumeration of walks, and it easily scales up to human (pan)genomes and transcriptomes. As it stores data in a modified FM-Index, GIN-TONIC may also be used for the compressed indexing of string-labelled graphs.

In particular, the data structure constructs a single modified FM-Index on a text encoding of the graph, allowing for an exact substring matching mechanism with quadratic complexity in the length of the query *Q* and number of graph vertices *V*, with a worst-case time complexity of *O*(|*Q*|^2^ |*V* |^2^ log |*V* | + *k*) in *O*(|*Q*||*V* | + *k*) space per query, where *k* is the number of occurrences of the query in the graph. This work also describes how such complexity can be further improved to *O*(|*Q*|^2^ |*V* | log |*V* | + *k*) time by approximating a solution to an NP-complete problem. The index itself can be constructed in *O*((|*V* | + |*E*|) log |*V* | + | Σ_*i*_ *S*_*i*_|) time, where Σ_*i*_ |*S*_*i*_| is the total length of the string labels and |*E*| is the number of edges. The space requirement depends on the FM-Index implementation and could range from compressed to expansive, but never exceeds *O*(|*V* | + | Σ_*i*_ *S*_*i*_|).

Finally, we describe an implementation of GIN-TONIC and provide experimental performance results in two biologically relevant use cases: human pangenome and transcriptome. Thanks to the addition of a suitable caching mechanism, GIN-TONIC is able to offer excellent performance despite having reasonable memory requirements. Code and benchmarks can be downloaded from https://github.com/uensalo/gin.

## 2. Methods

### 2.1. Notations

#### Strings

A string of length *L* over the finite alphabet Σ is defined as a sequence of characters *S* = *s*_0_*s*_1_ …*s*_*L*−1_ with *s*_*i*_ ∈ Σ ∀*i* ∈ {0,*…, L* − 1}, where *L* = |*S*| is the length of the string. We denote the set of all strings over the alphabet Σ as Σ^*\**^, the set of all strings of length *L* as Σ^*L*^, and the set of all strings up to length *L* as Σ^*≤L*^. Strings are indexed starting from 0, and the *j*^*th*^ character of a string *S* is denoted as *S*[*j* − 1]. The rank operation over a string *S* is defined as 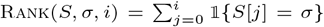 where *σ* ∈ Σ, and 0 ≤ *i <* |*S*|. A substring of a string *S* over the half open character range [*a, b*) is denoted as *S*[*a* : *b*] and defined as *s*_*a*_… *s*_*b*−1_ for *b > a*, |*S*| *> a >* 0, and |*S*| ≥ *b >* 0; it is taken to be *S*[*a* : *b*] = *ϵ* otherwise. ⊕ denotes iterative string concatenation, and defined to be 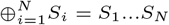 similar to the summation operator. ℬ (*S*) is the Burrows-Wheeler transform of *S*.

#### FM-Index lookup

Given some string query *Q*, the FM-Index of a string *S* returns the location of all indices *i*_0_, …, *i*_*m−1*_ such that *Q* occurs as a substring of *S* at *i*_*j*_ for *j* ∈ {0,*…, m*−1} through variations of Algorithm 1. The subroutine Advance-Range takes as input a half-open range over the suffix array and uses the *LF*-mapping to get a progressively refined range by walking through all the characters in the query.

#### String graphs

A string graph *G*(*V, E, 𝕊*) is defined as a set of vertices *V* = {*V*_1_,*…, V_N_*} labelled by non-empty strings 𝕊 = {*S*_1_,*…, S_N_*} such that *S*_*i*_ is the label of *V*_*i*_, and a set of directed edges *E* = {(*V*_*i*_, *V*_*j*_) | *V*_*i*_, *V*_*j*_ ∈ *V*}. The set of incoming neighbors for a vertex *V*_*i*_ is denoted as 𝒩 ^−^(*i*). A string

##### Algorithm 1

FM-Index Lookup

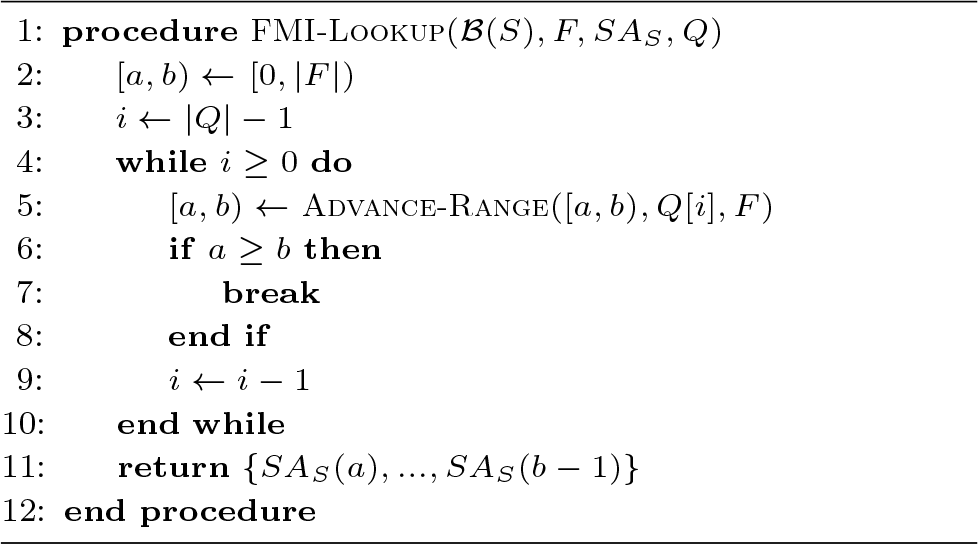

*S* is said to be induced or encoded by a string graph *G*(*V, E, 𝕊*) if there exists a walk 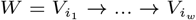 on *G* such that *S* is a substring of 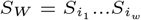. Conversely, the strings induced by a walk *W* are defined to be the set of all substrings of *S*_*W*_.

### 2.2. Problem Definition and Approach

The main problem of interest consists of enumerating all the starting points (“roots”) from which there exist walks inducing the given query. I.e.:

#### Definition 1

Let *G*(*V, E, 𝕊*) be a string graph, and let *Q* be a non-empty string over a non-empty alphabet Σ. We would like to enumerate all tuples (*V*_*i*_, *o*_*i*_), where *V*_*i*_ ∈ *V* and *o*_*i*_ ∈ ℕ, such that there exists a walk 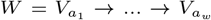 with the label concatenation 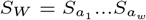 satisfying the following:

1. (The root of the walk is *V*_*i*_) 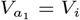.
2. (The walk induces *Q*) *Q* = *S*_*W*_ [*o*_*i*_ : *o*_*j*_] for some *o*_*j*_.
3. (The walk induces *Q* minimally) 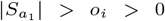 and 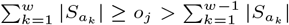.

We define three types of queries, which return:

1. Find: Suffix array ranges corresponding to walk roots
2. Locate: Walk roots in the graph domain as (*V*_*i*_, *o*_*i*_)
3. Walk: Reconstructed walks rooted at (*V*_*i*_, *o*_*i*_).

Locate queries first call Find queries to find suffix array ranges, and Walk queries call Locate queries to start traversing the graph from walk roots.

### 2.3. GIN-TONIC

This section defines the data structures used by GIN-TONIC.

**Definition 2 GIN-TONIC** is a data structure defined as a 4-tuple (I_FMI_(*G,* Π), 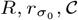) where:

- Π : ℕ → ℕ is a permutation over the set of vertices of a string graph *G*.
- I_FMI_ is an FM-Index constructed over a *graph encoding* GE(*G,* Π).
- 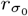 is a lookup table translating ranks of a special character *σ*_0_ in the Burrows-Wheeler domain of the graph encoding GE(*G,* Π) to corresponding ranks in the text domain.
- *R* is a data structure mapping ranges of a special character *σ*_0_ to ranges of *σ*_1_ based on *E*, over SA_GE(*G,*Π)_.
- 𝒞 is an optional, pre-computed cache of suffix array ranges corresponding to all substrings of length 1*…d* induced by the graph.

Definitions for GE(*G,* Π), 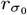, *R*, and *C*, follow.

### 2.4. The Graph Encoding

The graph encoding GE(*G,* Π) of a string graph *G* over a permutation Π is a string using special reserved characters to encode information on the permutation Π, the string labels *S*_*i*_ ∈ 𝕊 of *G*, and the mapping between the indices of vertices *V* of *G* and the string labels of *G*. The query algorithm will use the special characters to jump to other vertices when necessary.

**Definition 3** The **graph encoding of a string graph** is defined as the following string concatenation:

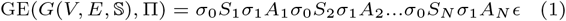

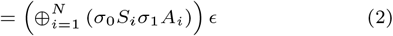

where Π denotes some permutation from {1,*…, N*} to 𝕊 {1,*…, N*}, *S*_*i*_ ∈ ? are strings over the string label alphabet Σ; *σ*_0_,*σ*_1_ ∈*/* Σ are two special characters respectively indicating the start and end of the string label *S*_*i*_ associated with each vertex *V*_*i*_, and *A*_*i*_ are strings over another alphabet Σ_Π_ such that Σ ∩ Σ_Π_ = ∅ and *σ*_0_,*σ*_1_ ∈*/* Σ_Π_, and the lexical order of *A*_*i*_ depends on Π. Hence, the alphabet of string GE(*G,* Π) is Σ ∪ Σ_Π_ ∪ {*σ*_0_,*σ*_1_}. The lexicographic ordering of the alphabet of GE(*G,* Π) is defined as 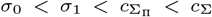 for any 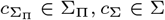,*c*_Σ_ ∈ Σ.

Positionally associating each vertex label with a *σ*_0_ and querying its rank allows one to obtain the index of the vertex in the text domain. When traversing the LF-mapping of the graph encoding, encountering a non-empty range of *σ*_0_ indicates that the current match has exhausted one or more vertex labels starting from which the currently matches suffix is induced, and the matching process might succeed by considering incoming neighbors if there are any. This property can be exploited to index all walks on the graph.

### 2.5. The Range Translator

The range translator *R* encodes the connectivity of *G* defined by *E* in the BW-domain. This is achieved by associating ranges of *σ*_0_, which express edge destinations, to ranges of *σ*_1_, which express edge sources. The query algorithm uses this mapping to traverse multiple vertices whenever necessary during matching.

#### Definition 4

Given two maps 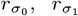 that can be computed from SA_GE(*G,*Π)_, with 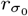 mapping ranks of *σ*_0_ from B(GE(*G,* Π)) to GE(*G,* Π), and 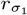 mapping the ranks of *σ*_1_ from GE(*G,* Π) to B(GE(*G,* Π)), the **single-vertex range translator** 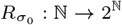 is defined as

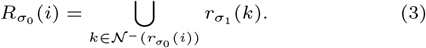

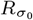 expresses as ranges of *σ*_1_ in SA_GE(*G,*Π)_ the notion of incoming neighbours of a vertex marked with a *σ*_0_. The **range translator** *R* : 𝕀→ 2^𝕀^, where 𝕀 = {[*a, b*]| *a, b* ∈ ℕ,*a* ≤ *b*}, is defined as

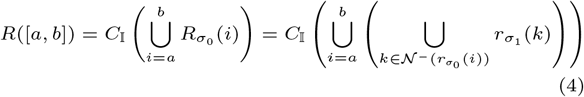

where *C*_𝕀_ : 2^ℕ^ → 2^𝕀^ is a function compacting a set of natural numbers into a set of intervals. *R* expresses the incoming neighbors of a range of vertices in SA_GE(*G,*Π)_.

We implement *R* in GIN-TONIC as a binary tree where each node contains a range key [*a, b*] and a list of intervals as values, called the interval-merge tree (IMT). *R* can be computed in Θ(|*V* | log |*V* |) time with *O*(|*V* |) space through this tree, which is defined as follows:

1. Each node (denoted with *T*) is associated with two children nodes, one range, and an interval list.
2. A parent node with a range [*a, b*] splits its range between its two children: the left child has the key-range 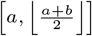, the right child has 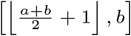.
3. Leaf nodes have key range [*k, k*], and interval list 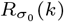).
4. A parent with children interval lists *ν*_*l*_, *ν*_*r*_ has a merged and compacted list of *ν*_*l*_ and *ν*_*r*_.

*R* can be evaluated on [*a, b*] by querying the tree recursively, as per Algorithm 2. Alternatively, implementations can merge references from all lists using a priority queue for a k-way merge.

#### Algorithm 2

Interval Merge Tree Lookup

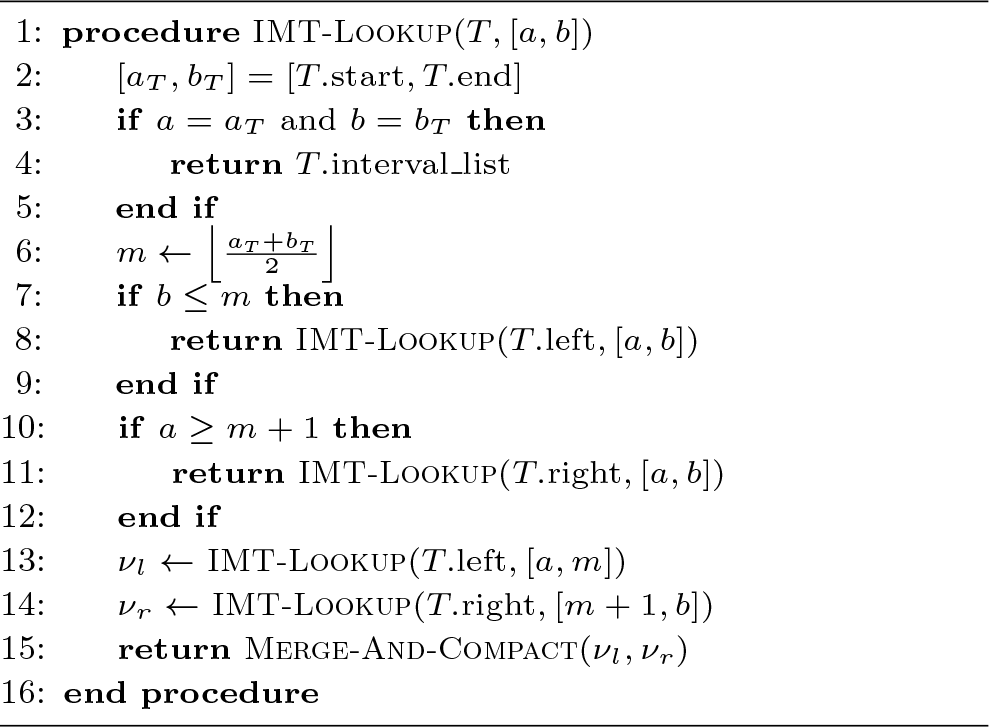

### 2.6. The Permutation

The permutation Π in GE(*G,* Π) is used to generate a string family *A*_*i*_ where each *A*_*i*_ follows each *σ*_1_ tied to vertex *i* according to Equation 1. This permutation dictates the order of *σ*_1_s in ℬ (GE(*G,* Π)); it allows to permute *σ*_1_s independent of the encoding and, possibly, to make suffix array ranges of walks that share a prefix consecutive, which increases the efficiency of querying. Π can be enforced by selecting a unique alphabet Σ_Π_, distinct from Σ ∪ {*σ*_0_,*σ*_1_}, generating *N* = |*V* | strings of length 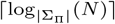, and arranging them lexicographically. Denote these sorted strings as *X*_1_,*…, X_N_*. Given any *σ*_1_, appending an *X*_*i*_ to it forces its rank in B(GE(*G,* Π)) to be *i*. Hence, setting 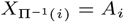 ensures that the *σ*_1_ with rank *i* in ℬGE(*G,* Π) has rank Π(*i*) in B(GE(*G,* Π)). Generally, using any sorted set of |Σ_Π_|-ary uniquely decodable codes for *X*_*i*_ achieves the same result with reduced overhead.

One can choose the permutation Π so that ranges of *σ*_1_s corresponding to the starting vertices of walks with a shared prefix are consecutive in the suffix array; that will result in faster query times. However, determining an optimally consecutive permutation is an NP-complete problem. Define

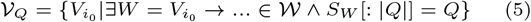

as the root of walks containing *Q*. To minimize interval counts returned by *R* for encountered ranges of *σ*_0_s during a query *Q*, define a constraint set *C*_*Q*_ for a query as

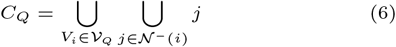

Choosing Π for all possible *Q* over all *C*_*Q*_s is equivalent to solving the Consecutive-Block-Minimization problem on a binary matrix *B* of size Σ^|Q|^ × |*V* | where each row corresponds to a query string *Q*, and for every *Q*_*i*_ at row *i*, the bit *B*[*i*][*j*] is set for all 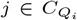. That yields an optimal Π, at the price of solving an NP-Complete but 1.5-approximable problem (34).

### 2.7. Querying and Construction Algorithms

This section provides two algorithms to query and construct GIN-TONIC, based on the definitions and observations provided and discussed in Section 2.3.

Algorithm 3 defines the construction of an GIN-TONIC from a string graph *G*, a permutation Π, and its associated *X*_*i*_. The procedure initially generates the graph encoding, calculates its suffix array, and sorts the sub-array of suffix rank-to-offset mappings for *σ*_0_s to derive the 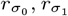. It then constructs an FM-Index using the suffix array. The overall computational complexity is *O*((|*V* | + |*E*|) log |*V* | + _*i*_ |*S*_*i*_|).

#### Algorithm 3 GIN-TONIC Construction

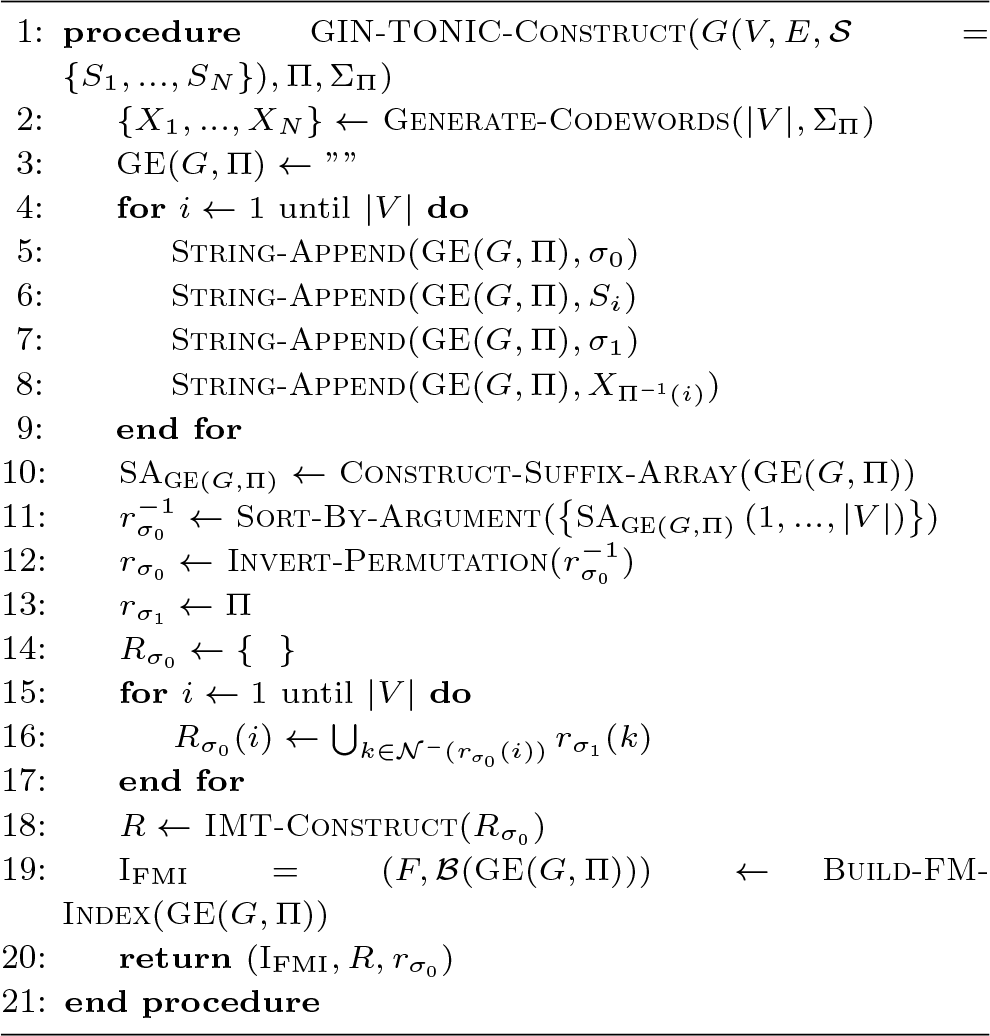

Algorithm 4 implements Locate queries, and returning *S* at line 27 instead of the decoded SA implements Find queries. Initially, the algorithm matches a query character to the graph encoding by updating the suffix array range based on the last character of the query over the full LF-mapping of GE(*G,* Π) in lines 2 − 4. This range then becomes the initial range to be tracked. Subsequently, the algorithm undergoes a loop, executing the ensuing steps for |*Q*| − 1 iterations:

1. Forking Phase (lines 6 − 16): For each tracked range, the algorithm assesses if the character *σ*_0_ is a preceding character in the LF-mapping of GE(*G,* Π). If so, then some walks must contain the ongoing partial match as a prefix (verified in lines 8 − 9). The algorithm then fetches the ranges of incoming neighbors using the range translator *R*. Each range from *R* is termed a “fork”, and such forks are collected in a separate set *F* (line 12).
2. Compaction Phase (line 16): The forks from *R* represent suffix array ranges over *σ*_1_. Although many forks might be retrieved, they can be compacted into fewer intervals. This phase determines the union of these intervals and simplifies their representation using the subroutine Compact-Intervals. Assuming *R* returns sorted ranges, this operation is linear in the number of intervals.

#### Algorithm 4 GIN-TONIC Lookup

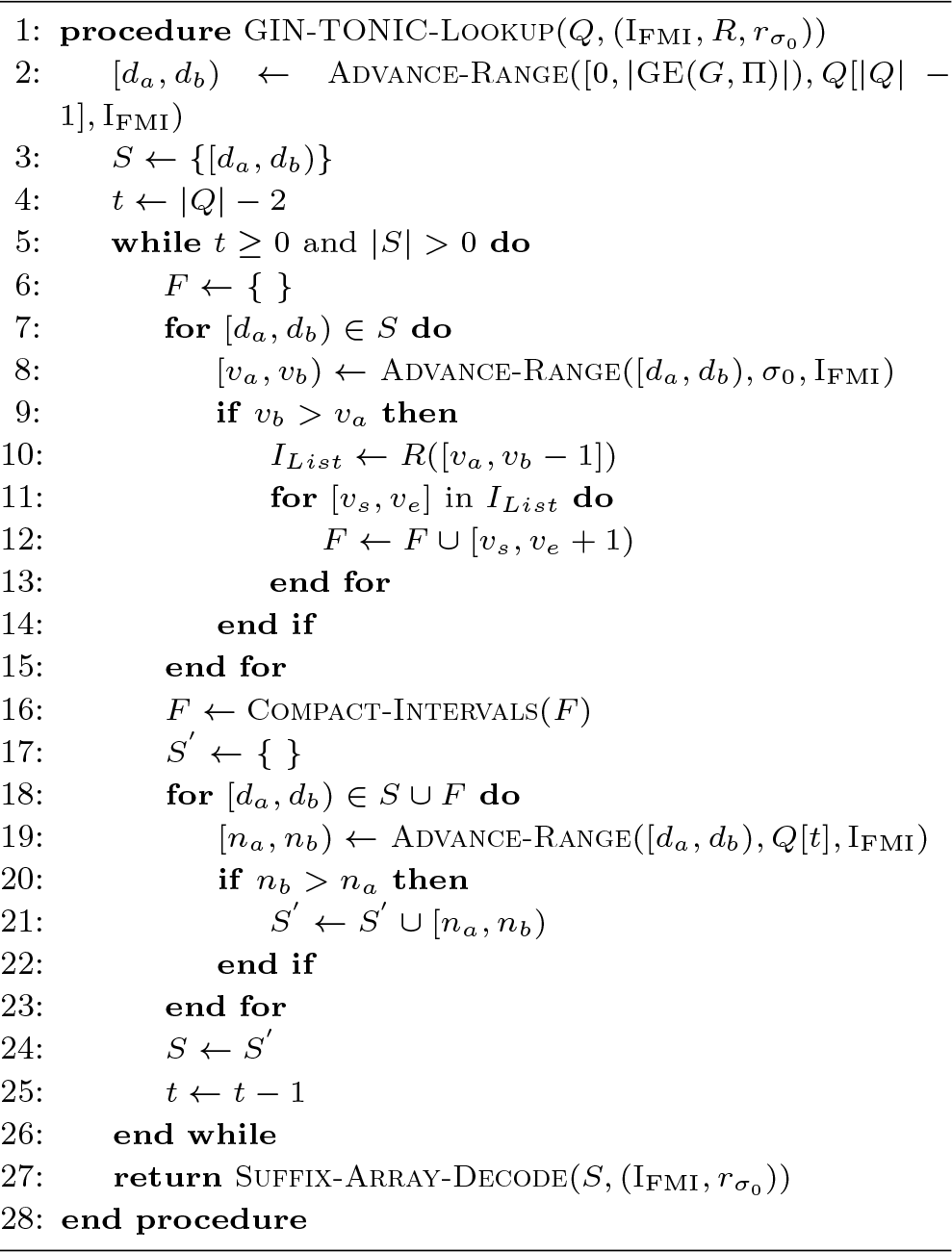
3. Advance Phase (lines 17 − 25): For each tracked range and fork, the algorithm advances to the next query character through the Advance-Range procedure on the FM-Index. A separate set *S*^′^ filters out ranges without the next character’s occurrence in the LF-Mapping. The algorithm then updates its tracked ranges to *S*^′^ and decrements the counter *t*, which indicates the next character position in *Q* to be considered.

After loop termination, *S* contains the suffix array ranges that represent the starting points of walks inducing *Q*. These results are converted to the text domain using Suffix-Array-Lookup. The algorithm’s time complexity is determined by the evaluation frequency of *R*, Advance-Range, and Compact-Intervals, and the range count returned by *R*. It can be demonstrated that at iteration *t*, the algorithm holds at most (*t* − 1)⌈|*V* |*/*2⌉ intervals in the worst case. This is because Compact-Intervals never returns more than ⌈|*V* |*/*2⌉ intervals by the pigeonhole principle. This leads to a worst-case time complexity of *O*(|*Q*|^2^ |*V* |^2^ log |*V* | + *k*), where *k* is the number of walk roots from which the query can be reconstructed. However, if Π is optimal, producing a single run in *B* for every prefix, then Compact-Intervals outputs the fewest possible intervals. This limits the intervals in *S* at iteration *t* to *t*. Consequently, the time complexity is reduced to *O*(|*Q*|^2^ |*V* | log |*V* | + *k*), as *R* is now called only *t* times rather than (*t* − 1)⌈|*V* |*/*2⌉. However, this bound is pessimistic unless GE is adversarially constructed in order to maximise the number of intervals returned by *R*. Typically, the time complexity depends on the number of occurrences in the graph of walk roots corresponding to the query.

### 2.8. The Pre-computed Suffix Array Range Cache

In general, the initial iterations of a query yield disjoint suffix array ranges returned through *R*, most of which are subsequently discarded during the algorithm’s advance phase, leading to unnecessary computation. A suffix array cache 𝒞 with depth *d* is defined as a function 𝒞: Σ^*≤d*^ → 2^𝕀^ which takes a string argument *Q* with length at most *d* and returns the suffix array ranges containing walks inducing *Q*. By looking up 𝒞 and initializing *S* at line 3 of Algorithm 4, then iterating only *t* − *d* times, we can significantly cut down redundant computations from earlier iterations, albeit with increased storage demands. While one could represent 𝒞 with a hash table, its static nature allows for a compressed FM-Index implementation. Given unique string keys *S*_1_,*…, S_K_* and their related suffix array interval lists *I*_1_,*…, I_K_*, we can define the cache encoding as 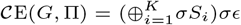, where *σ* ∈*/* Σ is a unique marker. In 𝒞E, the rank of *σ* matches the string key index. The interval lists are indexed using an FM-Index over 𝒞E, and the lists are permuted as 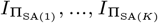, with Π_SA(*i*)_ indicating the permutation of *σ*s in ℬ (𝒞 E). To look up this index for a string *Q*, one matches *σQσ* against the FM-Index and navigates to the interval list 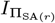, where *r* is the rank of *σ* after matching *σQσ*. Due to its structure, the FM-Index will have at most one match for *σQσ*. 𝒞 enhances query efficiency considerably.

## 3. Experiments and Results

### 3.1. Benchmark Setup

Performance of the proposed algorithms was assessed on two use cases. The first one involves a human pangenome graph, constructed from 20 haplotype-collapsed assemblies as described in (13), with 148,618 vertices, 214,995 edges, and a total label length of 3.14 × 10^9^ base pairs. The second use case considers the splicing graph of a human transcriptome with 1,029,466 vertices, 1,523,321 edges, and a total label length of 6.5 × 10^9^ base pairs. This was generated by overlaying the annotations of transcripts from GENCODE Release 40 (35) onto the human reference genome GRCh38.p13, with forward and reverse strands considered separately in the same file. Additional statistics for the two graphs are presented in Table 1. Experiments were conducted utilising a single thread on a machine with an AMD Ryzen Threadripper 3990X running at 2.2 GHz, 256 GB DDR4 RAM clocked at 2.7 GHz, and a 1 TB SSD with maximum read/write speeds of 7.0/5.0 GB/s.

**Table 1.** Statistics for input items.

| Input Name | Pangenome | Transcriptome |
| --- | --- | --- |
| Size (MB) | 2,996.456 | 6,264.277 |
| \|V\| | 148,618 | 1,029,466 |
| \|E\| | 214,995 | 1,523,321 |
| Avg Label Length (bp) | 21,100 | 6,350 |
| Med Label Length (bp) | 265 | 161 |
| Std Label Length (bp) | 156,000 | 89,600 |
| Min Label Length (bp) | 1 | 1 |
| Max Label Length (bp) | 30,092,626 | 30,380,454 |
| Avg Indegree | 1.45 | 1.48 |
| Med Indegree | 1 | 1 |
| Std Indegree | 0.692 | 0.937 |
| Min Indegree | 1 | 1 |
| Max Indegree | 11 | 93 |
| Avg Outdegree | 1.45 | 1.48 |
| Med Outdegree | 1 | 1 |
| Std Outdegree | 0.689 | 0.873 |
| Min Outdegree | 1 | 1 |
| Max Outdegree | 8 | 71 |

For each input item, optimal permutations Π over the suffix array as described in Section 2.6 were approximated for all walk prefixes of length 12 through simulated annealing; GIN-TONICs were then generated with rank and suffix array sampling periods of 32. Construction times for all implementations including the SDSL-based ones, and resulting index sizes, are listed in Table 2. They are comparable to those of a linear FM-Index, as the length of GE is approximately equal to the length of the human genome. To evaluate querying and decoding efficiency, 65,536 adversarial test queries maximising the number of spanned vertices were sampled from the input file for each one of lengths 2^4^,*…,* 2^12^. In addition, caches for each index with depths ranging from 1 to 12 were built as described in Section 2.8 and summarised in Table 3. Then, both our native FM-Index implementation and two different implementations based on the sdsl (36) (Huffman-shaped wavelet-trees with RRR bitvectors (11) and with hybrid bitvectors (37)) were benchmarked. Detailed logs, along with the relevant scripts used for data collection, are available at https://github.com/uensalo/gin/tree/master/benchmark.

**Table 2.**
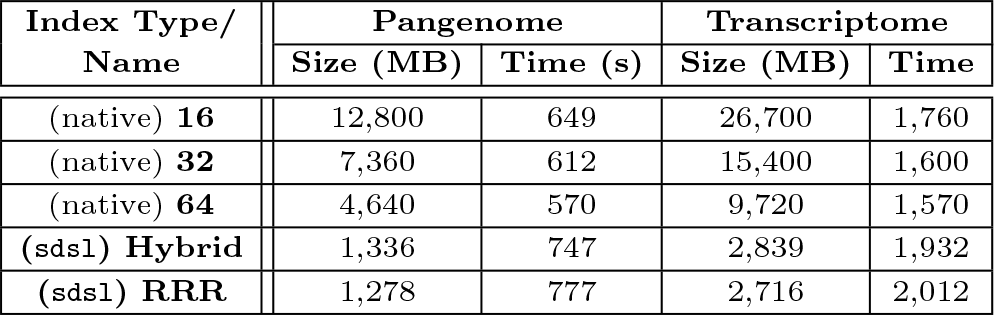
Index construction times, and sizes. (native) refers to our implementation, and (sdsl) to a compressed implementation using Huffman-shaped wavelet-trees on top of a particular bit-vector.

**Table 3.**
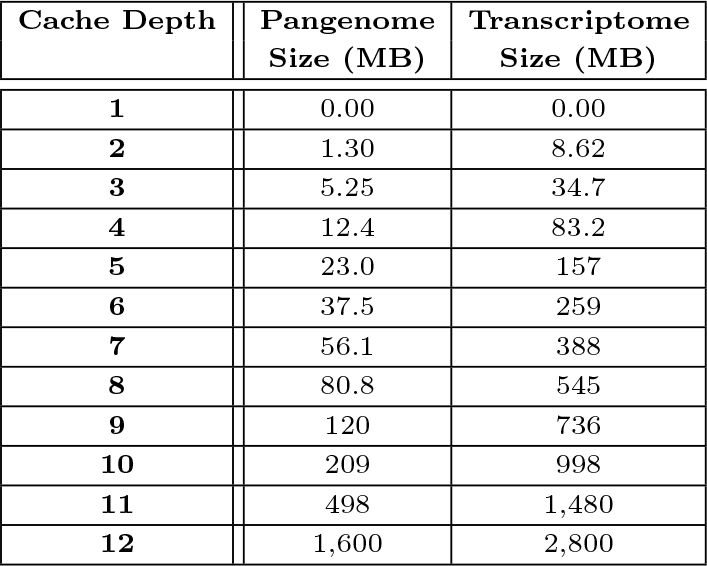
Cache sizes as a function of depth for both input items.

| Cache Depth | Pangenome<br>Size (MB) | Transcriptome<br>Size (MB) |
| --- | --- | --- |
| 1 | 0.00 | 0.00 |
| 2 | 1.30 | 8.62 |
| 3 | 5.25 | 34.7 |
| 4 | 12.4 | 83.2 |
| 5 | 23.0 | 157 |
| 6 | 37.5 | 259 |
| 7 | 56.1 | 388 |
| 8 | 80.8 | 545 |
| 9 | 120 | 736 |
| 10 | 209 | 998 |
| 11 | 498 | 1,480 |
| 12 | 1,600 | 2,800 |

**Table 4.** Total time taken in seconds for Find queries and walk root decodes, and total walk root counts (#) for 65,536 adversarial queries over the input items.

| Dataset | Pangenome |  |  | Transcriptome |  |  |
| --- | --- | --- | --- | --- | --- | --- |
| Query Length | Find (s) | Decode (s) | # | Find (s) | Decode (s) | # |
| 16 | 2.3 | 27 | 33,858,913 | 1.1 | 17 | 19,483,058 |
| 32 | 8.2 | 2.6 | 3,054,580 | 3.2 | 0.63 | 635,127 |
| 64 | 12 | 0.44 | 256,545 | 6.2 | 0.19 | 68,855 |
| 128 | 19 | 0.28 | 99,662 | 13 | 0.19 | 66,082 |
| 256 | 27 | 0.25 | 78,184 | 24 | 0.18 | 65,562 |
| 512 | 47 | 0.22 | 75,429 | 47 | 0.18 | 65,537 |
| 1,024 | 92 | 0.25 | 75,517 | 94 | 0.23 | 65,537 |
| 2,048 | 180 | 0.28 | 75,517 | 180 | 0.24 | 65,536 |
| 4,096 | 350 | 0.33 | 75,517 | 350 | 0.29 | 65,536 |

### 3.2. Effective Query Length

While the FM-Index traverses the LF-mapping exactly as many times as the length of the query *Q*, Algorithm 4 implies additional traversals every time the query needs to progress through the edges of the graph. This notion is captured by defining the *effective query length* |*Q*|^′^ as the number of LF-mappings done before the algorithm terminates, i.e., as the number of invocations of Advance-Range on line 20 of Algorithm 4. To quantify the overhead against the FM-Index, we also define the *relative effective query length* as

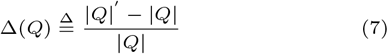

Δ(*Q*) will be 1 for a linear genome and *>* 1 for a graph genome, its precise value being fundamentally determined by the structure and complexity of the graph. Notably, Δ(*Q*) is a pure factor independent of the actual FM-Index used by GIN-TONIC, and hence it is the same across all the implementations tested while benchmarking GIN-TONIC. However, Δ(*Q*) can be decreased by optimizing the permutation Π and/or increasing the cache depth *d*, which are explored in the following sections.

### 3.3. Baseline Benchmarks

The main results of benchmarking GIN-TONIC’s baseline performance across different configurations (without optimisations; with optimised permutation; with cache; with both optimised permutation and cache) are shown in Figures 3a and 3b. Data for the cache & permutation case is reported for 3 different FM-Index implementations (native, sdsl with RRR bitvectors and sdsl with hybrid bitvectors).

**Figure 1.**
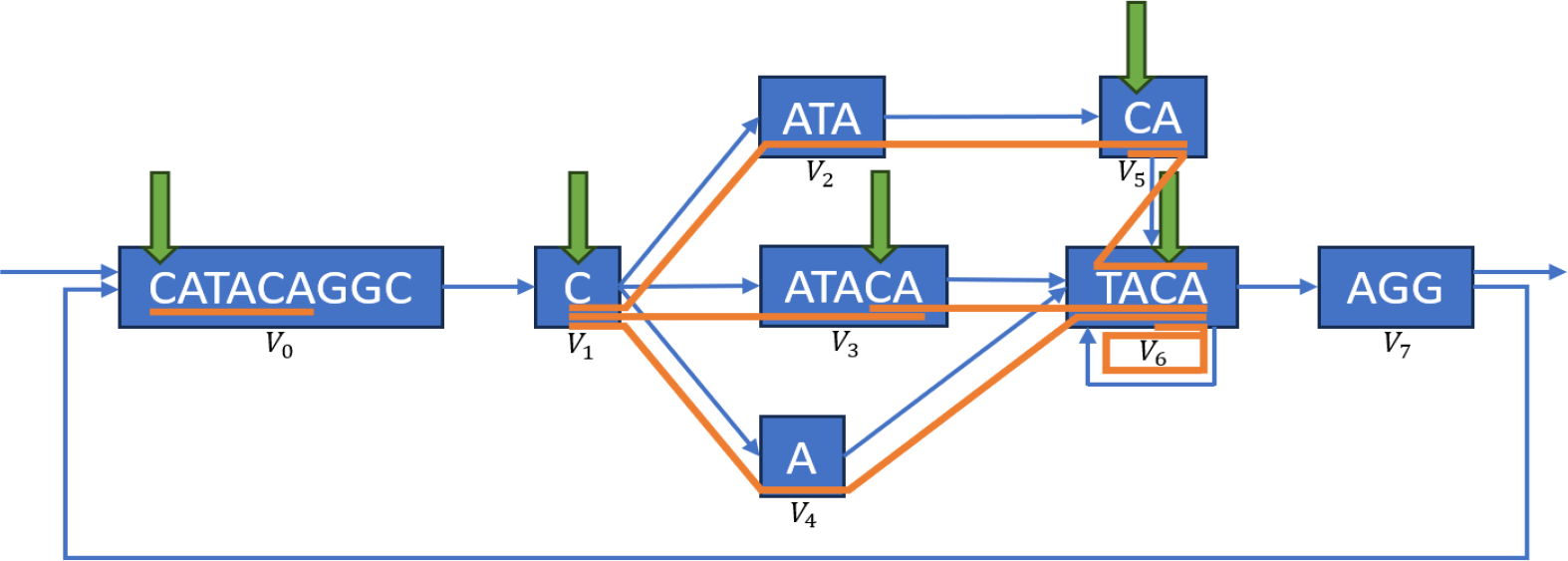
Example segment of a complex region of a string graph. Vertices are denoted with rectangles, edges with arrows, and vertex labels as text inside rectangles representing vertices. Walk roots corresponding to the string CATACA - equivalent to vertex index and offsets - are marked with green arrows. The walks inducing CATACA starting from the marked roots are marked with orange lines. Note that one root may induce multiple walks of the example string, that walks stemming from different roots may partially overlap, and that the same path can contain more than one match. Locate queries output all green arrows, and Walk queries output all orange paths.

**Figure 2.**
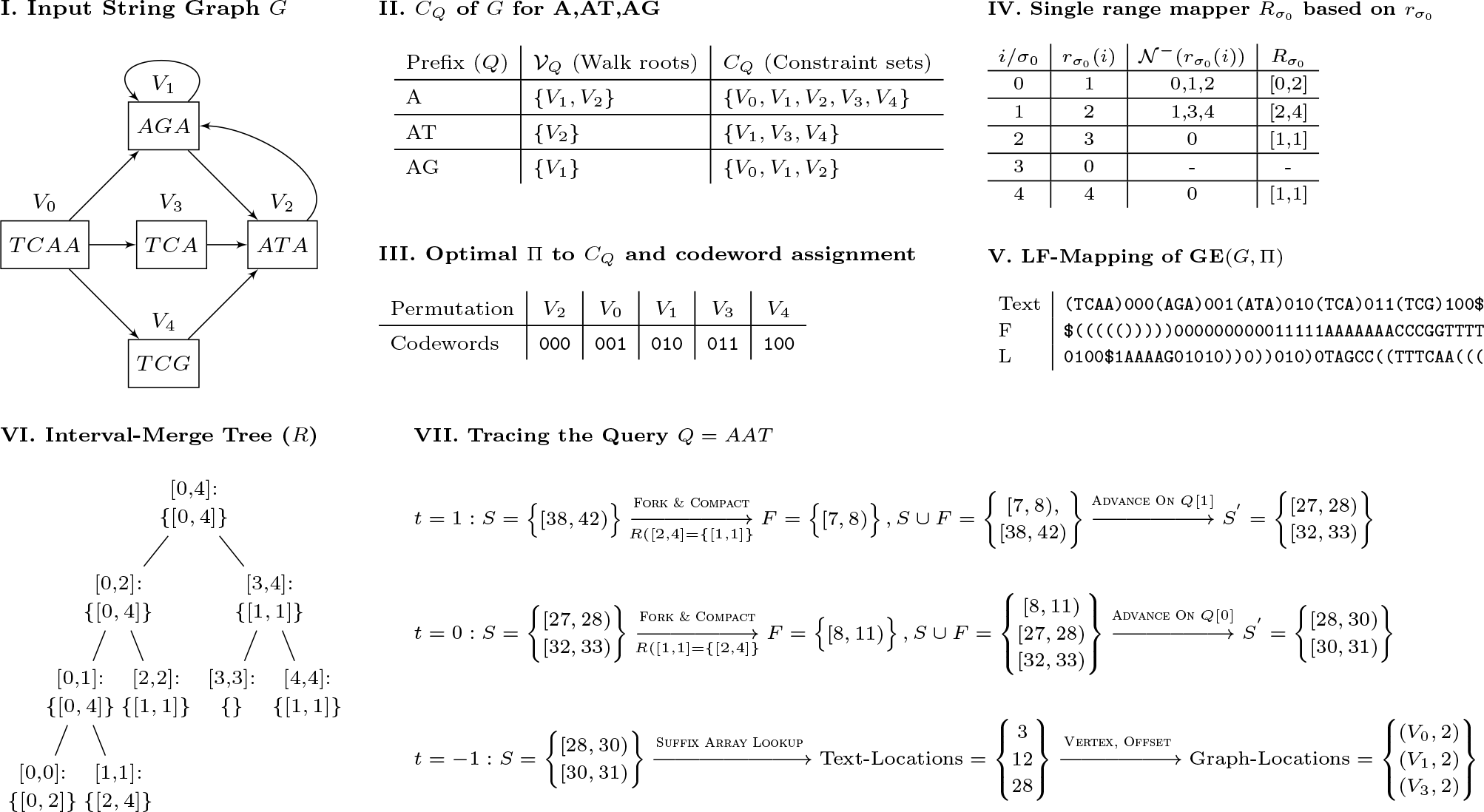
Illustrated examples of how to build and query a GIN-TONIC. **I.** A string graph of five vertices and 8 edges. **II.** Identified constraint sets for three prefixes *A, AT, AG*. **III.** Codeword assignment to enforce a permutation preserving a consecutive order of the constraint sets in II. **IV.** The single-vertex range translator over *σ*_0_. **V.** LF-Mapping constructed on the graph encoding with the permutation obtained in Step III. **VI.** Interval-merge tree based on the single range mapper 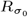. **VII.** Matching *Q* = *AAT* using the data structures in V and VI. The query is successively extended one character at a time starting from its end to the beginning as in the linear FM-Index.

**Figure 3.**
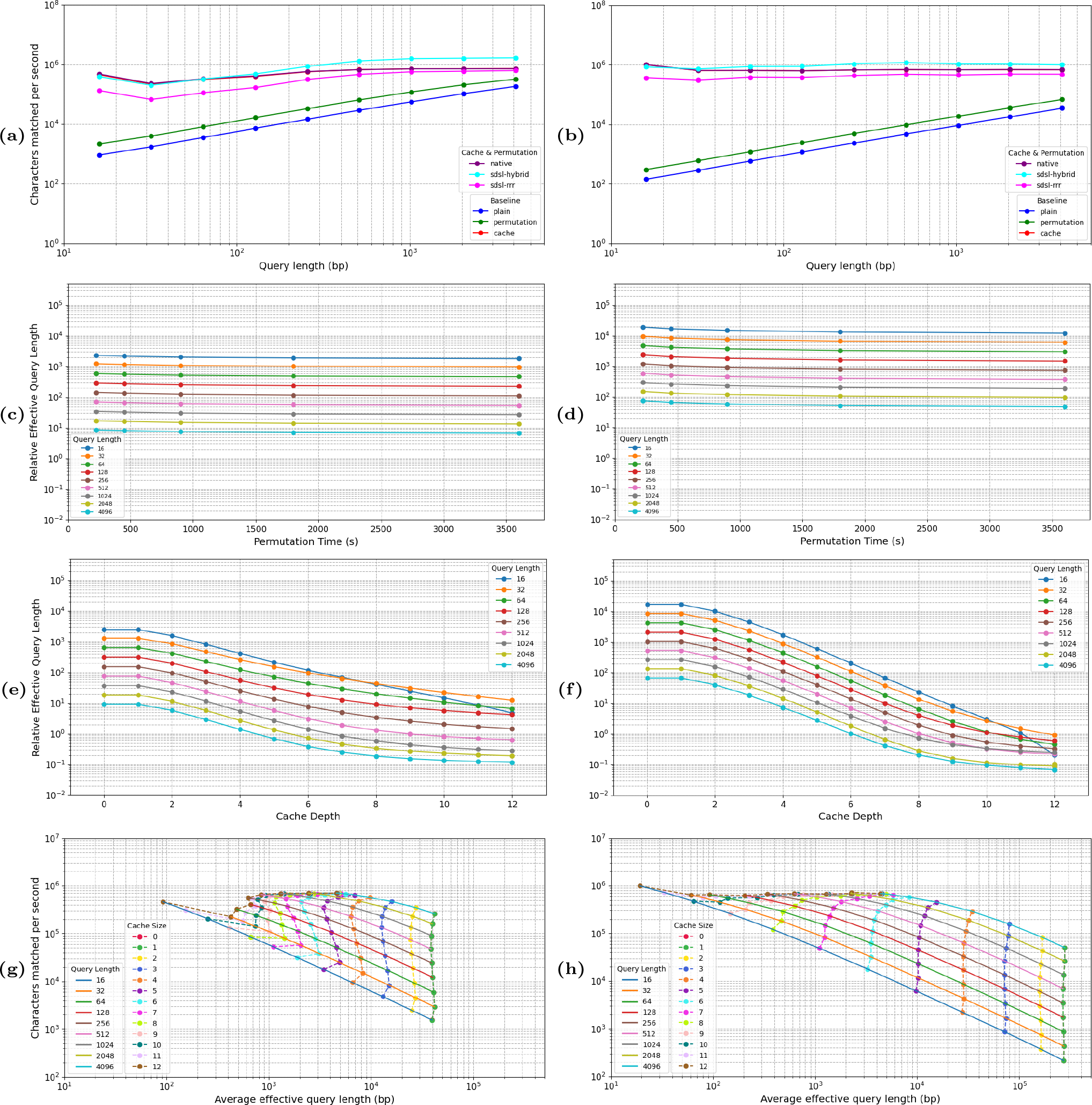
Benchmarking GIN-TONIC on adversarial queries. Left panels are for the pangenome, right panels for the transcriptome. Panels **(a,b)** show baseline performance using neither cache nor permutation, only permutation, only cache; and performance of different GIN-TONIC implementations (native and sdsl-based) using both cache and permutation as characters matched per second. Panels **(c,d)** show average Δ(*Q*) as a function of permutation optimization times. Panels **(e,f)** show average Δ(*Q*) as a function of cache depth. Caches of depth 0 and 1 have very similar performances close to the baseline since they cannot traverse more than one vertex, and hence they appear superimposed in the plots. Panels **(g,h)** show Δ(*Q*) vs. characters matched per second, grouped by cache depth (dashed lines) and average effective query length (solid lines).

Caches significantly outperform permutations in enhancing GIN-TONIC’s speed. An optimized permutation roughly doubled the effective query length and query throughput, whereas a cache led to several orders of magnitude of improvement (3 to 4 for short queries). However, using a permutation in addition to the cache can sometimes increase performance further; even though the transcriptome saw negligible gains, the pangenome showed around a 10% improvement for shorter queries.

In addition, the use of a cache effectively compensates for the complexity injected into the FM-Index by the graph structure. While without cache the number of characters matched per second by GIN-TONIC is not constant, unlike what happens with the FM-Index, that is not the case when a cache is used.

As for the effects of the implementation, the best performance is obtained from sdsl with hybrid bitvectors and the worst from sdsl with RRR bitvectors; while the behaviour of our native implementation is intermediate, being as fast as sdsl-hybrid at short query lengths and as slow as sdsl-RRR at longer queries. GIN-TONIC based on sdsl-hybrid reaches ∼10^6^ characters matched per second, which is comparable with what is achieved by an FM-Index on linear text.

### 3.4. Effect of Permutation

Figures 3c and 3d show the relation between the time spent optimising the permutation and the relative effective query length Δ(*Q*), which measures the slowdown introduced by the topology of the graph with respect to the linear FM-Index. No cache was used here, as the speed-ups provided by the two mechanisms are not independent. While the permutation can make the index more efficient, its influence is relatively modest with respect to that of the cache. It’s also worth noting that even though optimising the permutation does improve querying efficiency, it also incurs a one-off time cost at indexing time. In our case, we optimised the permutation for an hour at most, which is comparable to the time to build the FM-Index itself.

### 3.5. Effect of the Cache

Using a fixed permutation, cache depths from 1 to 12 were tested with rank and suffix array sampling rates set to 32; the results are illustrated in Figures 3e and 3f. They show that for both input graphs and for any given cache depth, Δ(*Q*) decreases as the query length increases; i.e., the longer the query, the closer the performance of GIN-TONIC was to that an FM-Index, and the more irrelevant the graph structure was. That can only happen if the number of jumps considered for longer queries decreases with the query length. We also observed that increasing the cache depth for a fixed query yielded several orders of magnitude of decrease in Δ(*Q*) for both input graphs, which indicates that a deeper cache is more able to optimise away the additional computation caused by the graph topology with respect to a linear FM-Index. Caches of order 10-12 make the values of Δ(*Q*) sufficiently close to 0 for GIN-TONIC to be practically usable.

### 3.6. Effect of Graph Structure

Strikingly, our benchmarks shine a light on the fundamental structural properties underlying the different graphs, as they directly determine the querying slowdown measured by Δ(*Q*).

Considering the actual performance of GIN-TONIC without a cache, one would expect the characters matched per second to be essentially independent of the query length starting from some query length on. After matching the first characters in the query, extending the query becomes cheap, thus rendering the work needed to match the rest of the query negligible. Hence, the expected curve showing Δ(*Q*) vs. characters matched per second of GIN-TONIC without a cache over queries of increasing lengths is vertical. That can indeed be observed for both input graphs in Figures 3g and 3h. Note that points associated with caches of depth 0 and 1 coincide as they offer very similar, negligible improvements over the baseline.

For both the pangenome and transcriptome, increasing cache depth resulted in the transformation from a vertical curve, meaning that Δ(*Q*) was independent of the query length, to a horizontal curve, which means that the high cost for extending the first few characters in the query is eliminated by the cache, and the cost of extending the query by one character tends to become independent of the query length. Above certain cache depths, the number of characters matched per second starts to stabilize, implying that other operations of the algorithm have become bottleneck.

However, there are differences between the pangenome and the transcriptome, as increasing the cache depth yields a flat curve for the transcriptome but not for the pangenome. This difference can be attributed to local graph structure. Vertices with shorter labels, high indegree and outdegree have the greatest potential to increase Δ(*Q*), as they will trigger more vertex traversals while extending the query. On the contrary, having a cache pre-computes the extensions of the query, and in the case where the label of a vertex is shorter than the depth of the cache, no additional computation arising from the topology of the graph is needed to extend the query.

This is confirmed experimentally by the local structure of the pangenome and the transcriptome. Figure 4 shows the distribution of vertex label length vs. vertex degree for each graph, and one can see that the distribution of vertex lengths in the pangenome is skewed towards longer lengths when compared to that for the transcriptome. In particular, a significant number of vertices of length ∼300 is present, which is probably an artefact pangenome construction; that is the likely reason why a flat curve is not achieved by a cache of depth 12. On the other hand, the vertex length distribution for the transcriptome is smooth and centered on much smaller values, which likely explains the effectiveness of the cache at decreasing Δ(*Q*).

**Figure 4.**
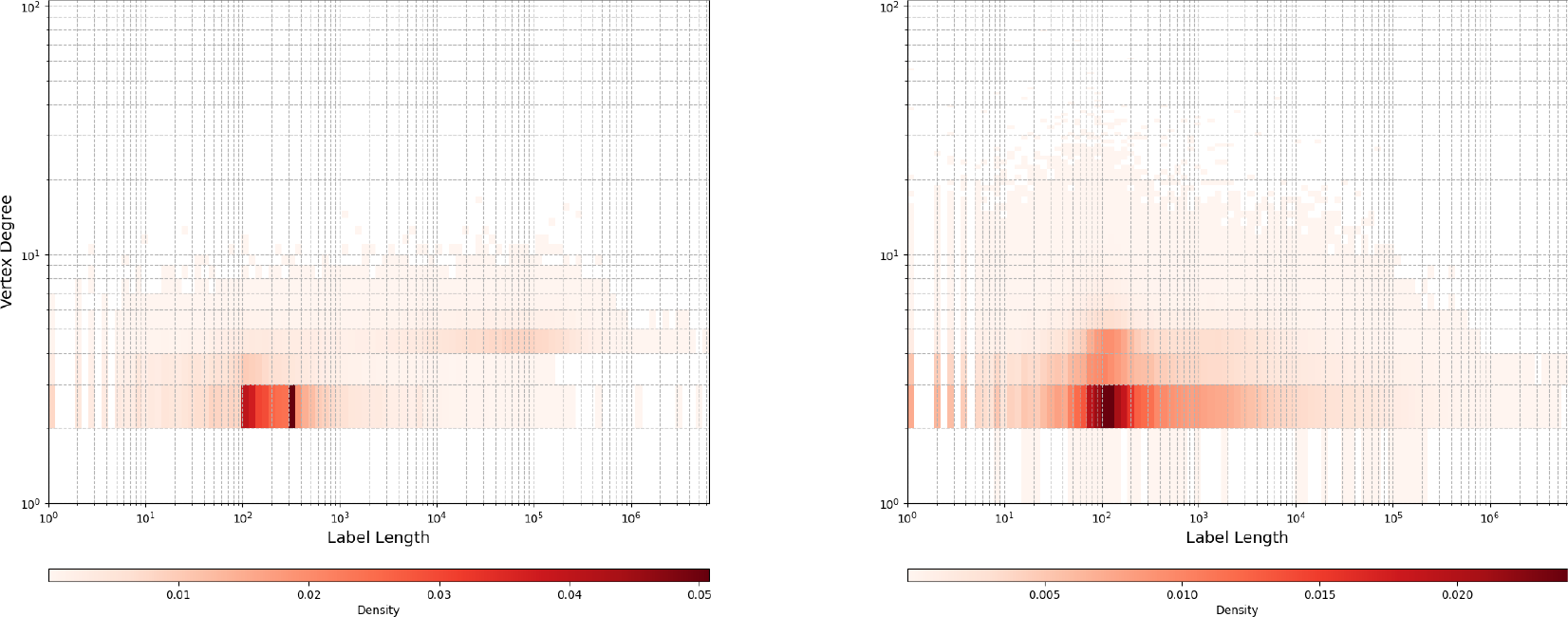
Vertex label lengths vs. degrees for each vertex for pangenome and transcriptome.

### 3.7. Locate and Walk Queries

Practical use of the index requires decoding suffix array ranges of matches *Q* into offsets of GE(*G,* Π). This decoding is independent of the internal details of Find queries, thus allowing query matching and suffix array range decoding into walk roots to be performed independently. In such a setting, the limiting factor in speed will be the slower of the two tasks.

Table 4 displays times spent performing Find queries of varying lengths and decoding the resulting suffix array ranges with the native implementation. For both inputs decoding was faster than Find for queries longer than 16 bp, thus making Find the bottleneck. However, for shorter queries, given the higher number of matches returned, decoding was slower.

After decoding suffix array ranges, the offsets in GE(*G,* Π) are not yet in the format (*V*_*i*_, *o*_*i*_). By using a prefix-summed array of the offsets at which vertices start in GE(*G,* Π), index and offset of the vertex can be found efficiently in *O*(log *V*) time via binary search. This operation is fast in practice and does not significantly affect decoding performance.

Walk queries cannot be answered via Algorithm 4; instead, a localised graph search can be used for each matching root (*V*_*i*_, *o*_*i*_) of *Q*, to reconstruct walks in the format 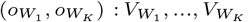, with offset 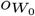 denoting the start of the match in 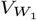 and offset 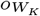 the end of the match in 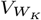. Graph labels can be bit-encoded to support Walk queries efficiently. This adds to the flexibility of GIN-TONIC, while also allowing the integration of other indexing strategies (e.g. haplotype indexing) to optimize the search space of walks.

## 4. Conclusions and Outlook

In this paper we have introduced GIN-TONIC, a monolithic data structure able to index arbitrary string graphs non-hierarchically. In terms of construction times, memory usage and querying and decoding times, its performance approaches that of an FM-Index over linear text, as demonstrated by experimental measurements of Δ(*Q*) on datasets of the size of a human pangenome and transcriptome. To the best of our knowledge, this work is the first one to achieve such a result.

The experimental performance reported in the paper could be improved further in several ways – for instance, by parallelising or pipelining several stages of the current implementation of GIN-TONIC, or by replacing its underlying FM-Index implementation with a hardware-optimised one. In addition, the representation of the string graph could be easily converted into a compressed and queryable file format and/or integrated into alignment software, possibly in conjunction with other haplotype indexing tools such as GBWT(20; 21). Future work will also explore how more flexible querying strategies taking into account biological priors could be integrated into GIN-TONIC.

## Contributions

**Ünsal Ö ztürk:** Conceptualisation, data curation, formal analysis, investigation, methodology, software, validation, visualisation, original draft, review, editing.

**Marco Mattavelli:** Conceptualisation, funding acquisition, project administration, resources, supervision, review, editing.

**Paolo Ribeca:** Conceptualistion, data curation, investigation, methodology, project administration, resources, software, supervision, validation, review, editing.

## Acknowledgements

Paolo Ribeca is affiliated to the National Institute for Health Research Health Protection Research Unit (NIHR HPRU) in Genomics and Enabling Data at University of Warwick in partnership with the UK Health Security Agency (UKHSA), in collaboration with University of Cambridge and Oxford. Paolo Ribeca is based at UKHSA. The views expressed are those of the authors and not necessarily those of the NIHR, the Department of Health and Social Care or UKHSA. A preliminary version of this work was presented in (38) as a poster.

## Data Availability

Instructions and all mentioned input data compatible with GIN-TONIC (39) are available freely on https://github.com/uensalo/gin-data. Files used to construct the transcriptome are available at https://www.gencodegenes.org/human/release_40.html (35). The human pangenome (GRCh38-20-0.10b) is available via (13).

## Notes

https://github.com/uensalo/gin

https://github.com/uensalo/gin-data

### Competing Interest Statement

The authors have declared no competing interest.

### Summary of Updates

Author affiliations were updated. An explicit availability and implementation statement was added just after the abstract. A data availability statement was added towards the end of the manuscript with a link to a repository containing the data used in the manuscript. A citation was added to an archived version of the software in the data availability section. One more citation (a survey on long read alignment) was added in the introduction section. The GitHub link to the benchmarks in Section 3.1 was updated to reflect the changes in the repository.

